# Signaling-based neural networks for cellular computation

**DOI:** 10.1101/2020.11.10.377077

**Authors:** Christian Cuba Samaniego, Andrew Moorman, Giulia Giordano, Elisa Franco

**Affiliations:** Department of Mechanical and Aerospace Engineering, University of California Los Angeles, California 90095, USA.; Department of Biological Engineering, Massachusetts Institute of Technology, Massachusetts 02139, USA.; Department of Industrial Engineering, University of Trento, 38123 Povo (TN), Italy.

## Abstract

Cellular signaling pathways are responsible for decision making that sustains life. Most signaling pathways include post-translational modification cycles, that process multiple inputs and are tightly interconnected. Here we consider a model for phosphorylation/dephosphorylation cycles, and we show that under some assumptions they can operate as molecular neurons or perceptrons, that generate sigmoidal-like activation functions by processing sums of inputs with positive and negative weights. We carry out a steady-state and structural stability analysis for single molecular perceptrons as well as for feedforward interconnections, concluding that interconnected phosphorylation/dephosphorylation cycles may work as multi-layer biomolecular neural networks (BNNs) with the capacity to perform a variety of computations. As an application, we design signaling networks that behave as linear and non-linear classifiers.

## I. Introduction

Living cells sense, process, and respond to a multitude of inputs from the environment, collectively using this information to make complex, life-sustaining decisions. To integrate and process several inputs at once, cells rely on signaling pathways that regulate downstream pathways depending on the input patterns they detect [1]. An important building block of signaling pathways is given by protein post-translational modification cycles, of which the best known is phosphorylation [2]. The rich dynamics and steady-state responses of these signaling pathway have been amply studied and are well-understood when they are taken as single-input, single-output isolated systems [3]. However, these pathways not only respond to multiple inputs, but they are also tightly interconnected, making it difficult to clearly identify their input-output function and their full potential for signal processing and computation.

In this manuscript, we consider a model for multi-input, single output phosphorylation/dephosphorylation cycles and describe their ability to operate as molecular perceptrons. Building on previous work by us and others, that proposed a strategy to build biomolecular neural networks (BNNs) using molecular sequestration [4], here we demonstrate that, under some assumptions, these natural signaling pathways can operate as multi-input perceptrons as long as their input-output behavior has a tunable threshold. Further, these molecular perceptrons can process several inputs assigning them positive and negative weights by tuning catalytic rate parameters; the presence of both positive and negative weights is essential for complex classification. We report a *structural* steady-state and stability analysis of the multi-input phosphorylation/dephosphorylation cycle model considered as a candidate perceptron, as well as stability analysis for BNNs resulting from the feedforward interconnection of multiple perceptrons. With simulations, we show that by tuning the weights suitably, and by designing the depth of the BNN, it is possible to build linear and non-linear classifiers.

There is a long history of theoretical and experimental work devoted to discovering the computational power of chemical and biological networks under the lens of neural networks. For example, a chemical implementation of neural networks based on reversible reaction was proposed in [5]; networks in living cells were examined in [6], showing that they present features not found in conventional computer-based neural networks; chemical Boltzmann machines have been described in [7]. It has been proposed that transcriptional networks *in vitro* and *in vivo* may operate as neural networks by tuning various reaction rate parameters [8], [9]. Nucleic acids have been engineered for experimental demonstrations of molecular neural networks and classifiers *in vitro* [10], [11], in cell extracts [12], and in cells [13], [11].

Although in practice any biological network may be tuned to work as a neural network [6], the main challenge is to translate an abstract design into an implementation with predictable behavior. A long-standing challenge, for example, is that of implementing negative weights in a biological neural network that can only admit positive variables (concentrations). In previous work, we demonstrated that molecular sequestration reaction can theoretically implement positive and negative weights because it performs a subtraction operation between its inputs [4]. Because a salient feature of molecular sequestration is that it generates an ultrasensitive response with a tunable threshold, we rationalize that other networks with a tunable threshold may implement molecular neural networks with positive and negative weights. This rationale motivates the present study: we show that when post-translational modification cycles, such as phosphorylation/dephosphorylation, present an ultrasensitive response with a tunable threshold, they can operate as perceptron nodes with positive and negative weights. After providing basic definitions in Section II, in Section III we report mathematical analysis of phosphorylation/dephosphorylation cycles as a perceptrons. In section IV we describe the design of a single-layer linear classifiers, Section V describes multilayer BNNs, and finally Section VI provides simulation examples of non-linear classifiers.

## II. Background: Artificial Neural Networks and Perceptrons

Artificial neural networks (ANNs), also called perceptron networks, take inspiration from the architecture of the human brain. ANNs consist of multiple, interconnected artificial neurons, or perceptrons, which represent the nodes of the network. Each perceptron performs a different computation of the same functional form, whose outcome depends on the perceptron’s inputs and parameters. We adopt the following model (Fig. 1A) for a single perceptron that takes *n* inputs:

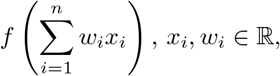

where *f* is a non-linear activation function, *x*_*i*_ are the perceptron inputs, *i* = 1*, …, n*, and *w*_*i*_ is the parameter or weight associated to the *i*th input. By connecting multiple perceptrons, each with potentially distinct inputs and weights, ANNs achieve the emergent ability to carry out complex computations far more sophisticated than those realizable by an individual perceptron [14]. In the next sections, we discuss how a well-known class of (post-translational) signaling networks can be viewed as molecular perceptrons.

**Fig. 1.**
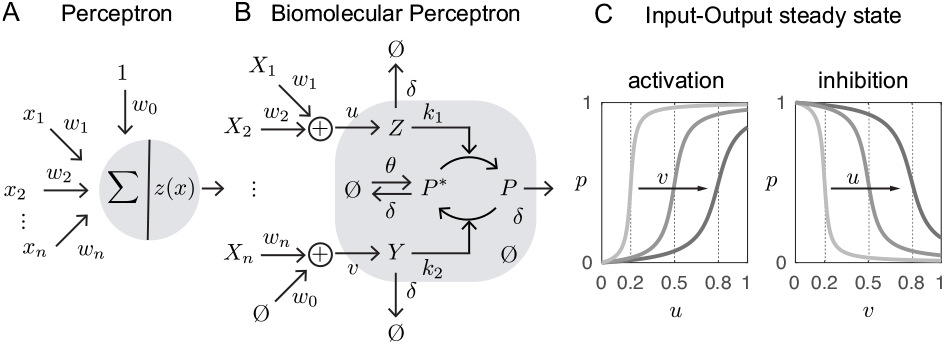
Phosphorylation-based perceptron. A: Perceptron scheme. B: Biomolecular implementation of the perceptron, including at its heart a phosphorylation-dephosphorylation cycle. C: Sigmoidal-like activation and inhibition functions.

## III. Phosphorylation-dephosphorylation cycles can operate like biomolecular perceptrons

Here we argue that post-translational modification cycles such as phosphorylation-dephosphorylation cycles can operate like biomolecular perceptrons (Fig. 1). Let us denote chemical species with uppercase letters, and their concentrations with the corresponding lowercase letter, so that species *X* has concentration *x*.

In a phosphorylation-dephosphorylation cycle (scheme in Fig 1B), a target protein *P*^*^ (inactive state, unphosphorylated) is converted into *P* (active state, phosphorylated) in the presence of a kinase *Z*, which adds a phosphate group to *P*^*^ so as to produce *P*. A phosphatase *Y* can reverse this process by removing the phosphate group from protein *P*, thus producing *P*^*^.

We model this process through the following biochemical reactions. The kinase *Z*, the phosphatase *Y* and the target protein *P*^*^ are produced at a constant rate, respectively *u*, *v* and *θ*:

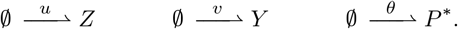

All species are subject to first order degradation with rate constant *δ*:

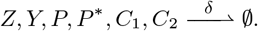

*Z* and *P*^*^ bind with rate constant *a*_1_ to form an intermediate complex *C*_1_, which dissociates at rate constant *d*_1_. The complex *C*_1_ produces *P* and *Z* with rate parameter *k*_1_.

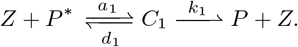

Analogously, *Y* and *P* bind at rate constant *a*_2_ to form an intermediate complex *C*_2_, which dissociates with parameter *d*_2_. In addition, the complex *C*_2_ can produce *P*^*^ and *Y* with rate parameter *k*_2_.

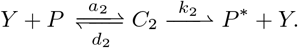

Assuming that the reaction kinetics follow the law of mass action, the dynamic evolution of the species concentrations can be expressed by the following system of Ordinary Differential Equations (ODEs):

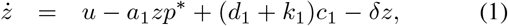

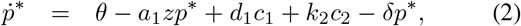

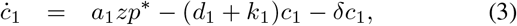

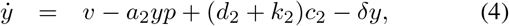

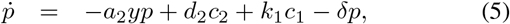

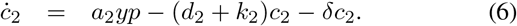

We can show that the system is positive, which is consistent with the fact that the state variables are concentrations of chemical species, hence cannot take negative values.

### Proposition 1

The system (1)–(6) is positive: given a positive initial condition, all the state variables have nonnegative values at all future times.

*Proof:* The result follows by noticing that, for each state variable *x ∈ {z, p*^*^*, c*_1_*, y, p, c*_2_}, 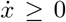 when *x* = 0, hence the variable cannot become negative.

### A. Stability analysis without production and degradation

If we neglect production and degradation reactions (*u* = *θ* = *v* = 0 and *δ* = 0), assuming that they compensate for each other, then the system simplifies to:

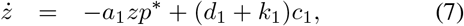

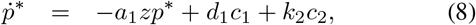

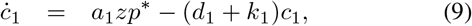

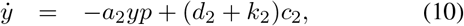

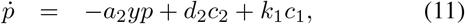

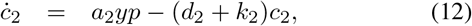

where the sums *z* + *c*_1_, *y* + *c*_2_ and *p*^*^ + *p* + *c*_1_ + *c*_2_ are constant quantities, hence the system can be associated with a reduced-order system with three differential equations and three conservation laws. Since each species is involved at least in one conservation law, this is a *conservative* chemical reaction network. The steady-state value and its global stability can only be assessed within the *stoichiometric compatibility class* associated with an assigned value of the conserved quantities.

#### Theorem 1

The system (7)–(12) admits a unique steady state within each stoichiometric compatibility class, and such a steady state is structurally globally asymptotically stable.

*Proof:* If we set *z* = *z*^*tot*^ − *c*_1_ and *y* = *y*^*tot*^ − *c*_2_, and neglect the dynamic equations for *z* and *y*, from (7)–(12) we get the reduced-order system

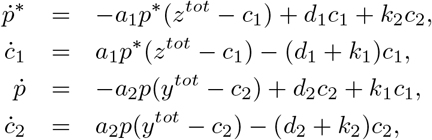

where *p*^*^ + *p* + *c*_1_ + *c*_2_ is constant. The Jacobian is

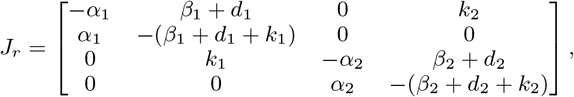

where 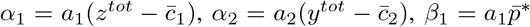 and 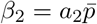. *J*_*r*_ is singular and weakly column diagonally dominant, with strictly negative diagonal entries and non-negative off-diagonal entries. Therefore, the system admits the 1-norm

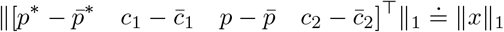

as a weak Lyapunov function that certifies the *structural* marginal stability of its steady state 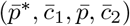 [15], [16, Section 4.5.5], regardless of the values of the system parameters. The only eigenvalue on the imaginary axis is *λ* = 0, associated with the left eigenvector [1, 1, 1, 1] that identifies an *invariant* subspace for the system. In this invariant subspace, the system admits a polyhedral Lyapunov function whose unit ball is given by the intersection between the shifted diamond centred at the steady state 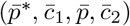 and the hyperplane orthogonal to the vector [1, 1, 1, 1] and passing through 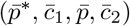. Since the system Jacobian is structurally nonsingular within the stoichiometric compatibility class, this Lyapunov function guarantees that the system equilibrium is *structurally* asymptotically stable. Note that, since the original (non-linearized) system can be rewritten as 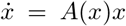, where *A* is related to the integral of the system Jacobian [15], [17], the system admits the same Lyapunov function at all points, hence a global (not only local) stability analysis can be conducted. In particular, in view of the structural nonsingularity of the Jacobian, the results in [17] also guarantee the uniqueness of the steady state and its *structural global asymptotic stability* within the stoichiometric compatibility class. See also [18], where the system (7)–(12) is shown to be *structurally attractive*.

It is worth stressing that Theorem 1 does not rely on the mass-action-kinetics assumption: the same result would hold true for any functional form (and parameter values) of the reaction kinetics, as long as the reaction rates are monotonic functions of the reagent concentrations.

### B. Quasi-steady-state approximation

In the presence of production and degradation reactions, the quantities *z*^*tot*^ = *z* + *c*_1_, *y*^*tot*^ = *y* + *c*_2_, and *p*^*tot*^ = *p*^*^ + *p* + *c*_1_ + *c*_2_ are not constant. From Equations (1)–(6), it follows that these quantities evolve over time as

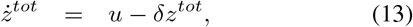

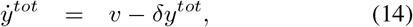

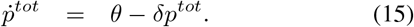

Therefore, at the steady state, 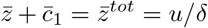, while 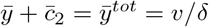 and 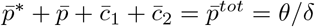.

Let us adopt a commonly accepted assumption in enzyme kinetics (cf. Michaelis-Menten kinetics) and consider that the concentrations of the intermediate complexes, *c*_1_ and *c*_2_, do not change on the time scale of product formation, thus reaching a quasi steady state:

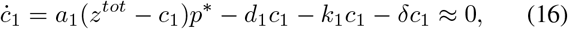

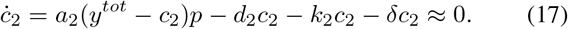

Therefore

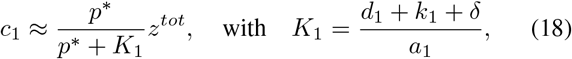

and

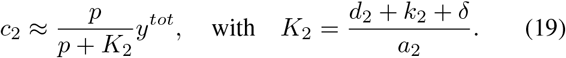

If we plug the expression of *c*_1_ and *c*_2_ into the equation for *ṗ*, we obtain

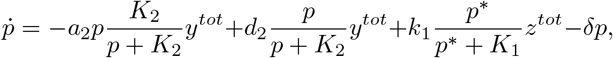

which can rewrite as

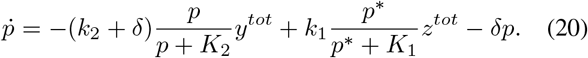

Similarly, the equation for *ṗ*^*^ can be rewritten as

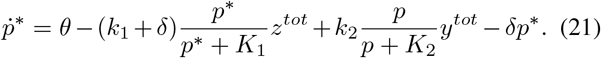

### C. Steady-state analysis

To perform the steady-state analysis of the phosphorylation-dephosphorylation network, we make the following simplifying assumptions, which are typically verified in practice. Phosphorylation and dephosphorylation are assumed to be much faster than degradation: *k*_1_*, k*_2_ ≫ *δ*. Also, the total amount of the target protein is assumed to be much larger than the total amount of the enzymes: *p*^*tot*^ ≫ *z*^*tot*^, *y*^*tot*^. Then, since *c*_1_ *< z*^*tot*^ ≪ *p*^*tot*^ and *c*_2_ *< y*^*tot*^ *p*^*tot*^, we can approximate *p*^*^ + *p* = *p*^*tot*^ − *c*_1_ − *c*_2_ ≈ *p*^*tot*^. Then, by considering *k*_1_ + *δ ≈ k*_1_ and *k*_2_ + *δ ≈ k*_2_ (since *k*_1_*, k*_2_ ≫ *δ*), we can rewrite equations (20) and (21) as

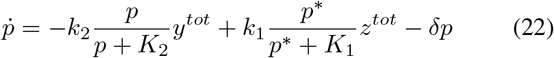

and

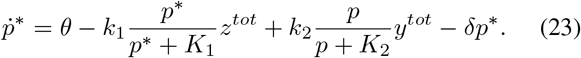

Under these approximations, *p* + *p*^*^ = *p*^*tot*^ and consistently *ṗ* + *ṗ*^*^ = *ṗ*^*tot*^.

Also, under the above approximations, by rearranging equation (22) at steady state (i.e., *ṗ* = 0) and neglecting the term *δp*, we can write the ratio

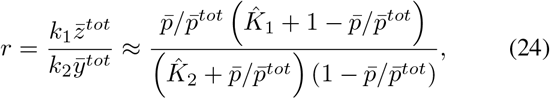

where 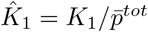 and 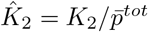.

In addition, to simplify our analysis, let us assume that 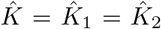. When the steady-state amount of active *p* is larger than half the total amount, 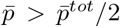, the ratio is larger than one: *r >* 1. Conversely, when 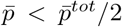, the ratio is smaller than one: *r <* 1. Finally, when 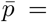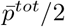, this results in *r* = 1 (the inputs 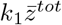 and 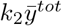 are approximately the same in magnitude). In short,

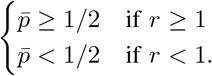

We can rewrite this expression as a function of 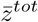 and 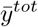 as follows:

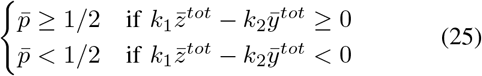

Since at steady state 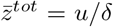 and 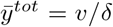, we can rewrite the condition as a function of the inputs *u* and *v*:

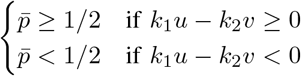

This mechanism allow us to implement both positive and negative weights, thanks to the fact that the input *u* appears in the condition with a positive sign, while the input *v* appears with a negative sign. Then, defining the input 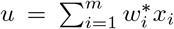, associated with a positive sign, and the input 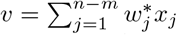, associated with a negative sign, the effective input (whose sign determines whether the value of 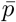 is above or below the threshold 1*/*2) becomes

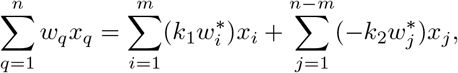

which corresponds to the weighted sum of inputs with both positive and negative coefficients. The coefficients 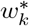 can be interpreted biologically as production rates of the enzymes *Z* and *Y*. Then, the steady-state expression for multiple inputs can be rewritten as

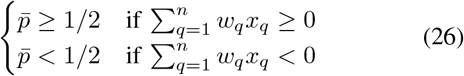

### D. Stability analysis

In this section, we show structural stability of the reduced order model including equations (13), (14) and (15), as well as equations (22) and (23). Since *p* + *p*^*^ = *p*^*tot*^, we can consider a system with only four equations:

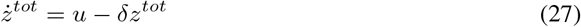

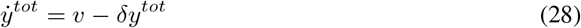

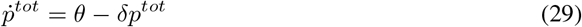

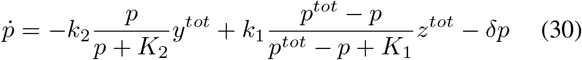

#### Theorem 2

The system (27)–(30) admits a unique steady state, which is structurally globally asymptotically stable.

*Proof:* We start by observing that the system trajectories are asymptotically bounded in the open set 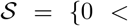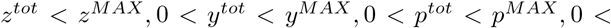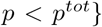, hence at least one steady state must exist in 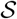. In particular, it can be seen that 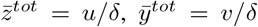 and 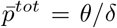, while 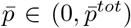 solves the equilibrium condition

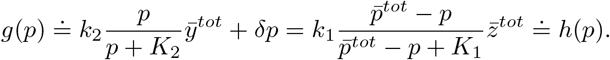

In the interval 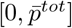, *g*(*p*) is a continuous strictly increasing function with *g*(0) = 0 and 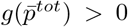, while *h*(*p*) is a continuous strictly decreasing function with *h*(0) *>* 0 and 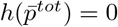, hence an intersection must exist in view of continuity, and it is unique in view of monotonicity.

Structural local asymptotic stability of the equilibrium can be proven by noticing that the Jacobian matrix

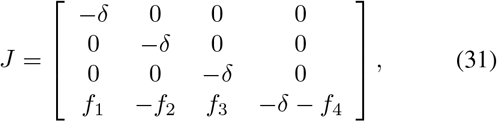

where the quantities 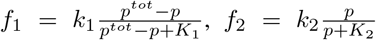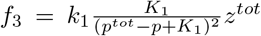 and 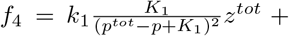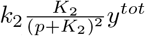 are positive, is a lower triangular matrix, whose eigenvalues are the negative diagonal entries.

Structural asymptotic stability can also be guaranteed by the existence of a polyhedral Lyapunov function [15]: the similarity transformation *TJT*^−1^, with *T* = diag(1, 1, 1*, ρ*), turns *J* into a column diagonally dominant matrix with strictly negative diagonal entries, if *ρ* is chosen small enough; then, the weighted 1-norm ||*x*||_1*,T*_ = ||*Tx*||_1_, with 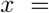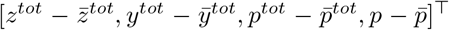, is a structural Lyapunov function for the system [16, Proposition 4.58]. Finally, since the Jacobian is structurally nonsingular, the results in [17] guarantee the *structural global asymptotic stability* of the equilibrium.

Also Theorem 2 would hold for any kinetics (not necessarily mass-action or Michaelis-Menten), as long as monotonicity with respect to species concentrations can be assumed.

#### Remark 1

Since all the variables of system (27)–(30) are asymptotically bounded in an open set, as shown in the proof of Theorem 2, the following fundamental result from degree theory can also be applied to prove uniqueness of the equilibrium.

#### Theorem 3

[19], [20] Assume that the system 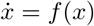, with *f*: ℝ*n* → ℝ*n* sufficiently regular, has solutions that are globally uniformly asymptotically bounded in an open set 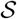 and admits *N <* ∞ equilibrium points 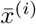, *i* = 1*, …, N*, each contained in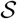 and such that the determinant of the system Jacobian matrix evaluated at 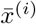 is nonzero: 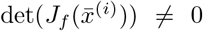 ∀*i*. Then, 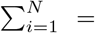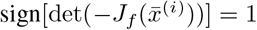.

For the Jacobian in Equation (31), det(−*J*) *>* 0 structurally; therefore, the equilibrium must be unique.

## IV. Tunable Molecular Linear Classifiers

In Section III, we have shown that a reaction network based on post-translational modification by phosphorylation and dephosphorylation may work as a tunable thresholding mechanism (see Fig. 1C). In particular, the threshold can be adjusted by the production of kinase and phosphatase species by positive and negative inputs, respectively, per equation (26). Altogether, this circuit can be viewed as a molecular perceptron with a sigmoidal activation function that incorporates positively- and negatively-weighted inputs.

In this section, we consider two case studies to show how the phosphorylation-dephosphorylation cycle, and by extension any other post-translational modification cycle, may be engineered to build bio-molecular linear classifiers in the presence of multiple inputs, and we discuss how adjusting the weights associated with kinetic rates can predictably program the linear classification.

### A. Network with two positive and one negative weights

We start with the design of a linear classifier with three inputs, as shown in Fig. 2A. We consider two inputs *X*_1_ and *X*_2_, which produce the kinase *Z* with rates *w*_1_ and *w*_2_, as well as a constant production rate *w*_0_ for the phosphatase *Y*. All other chemical reactions are taken as in Section III, as shown in Fig. 2B. Recall that, in our scheme, input species that produce *Z* are considered positively-weighted, while input species that produce *Y* are negatively-weighted. Thus, this reaction network represents a single perceptron with two positive inputs and a single, fixed negative input, or bias.

**Fig. 2.**
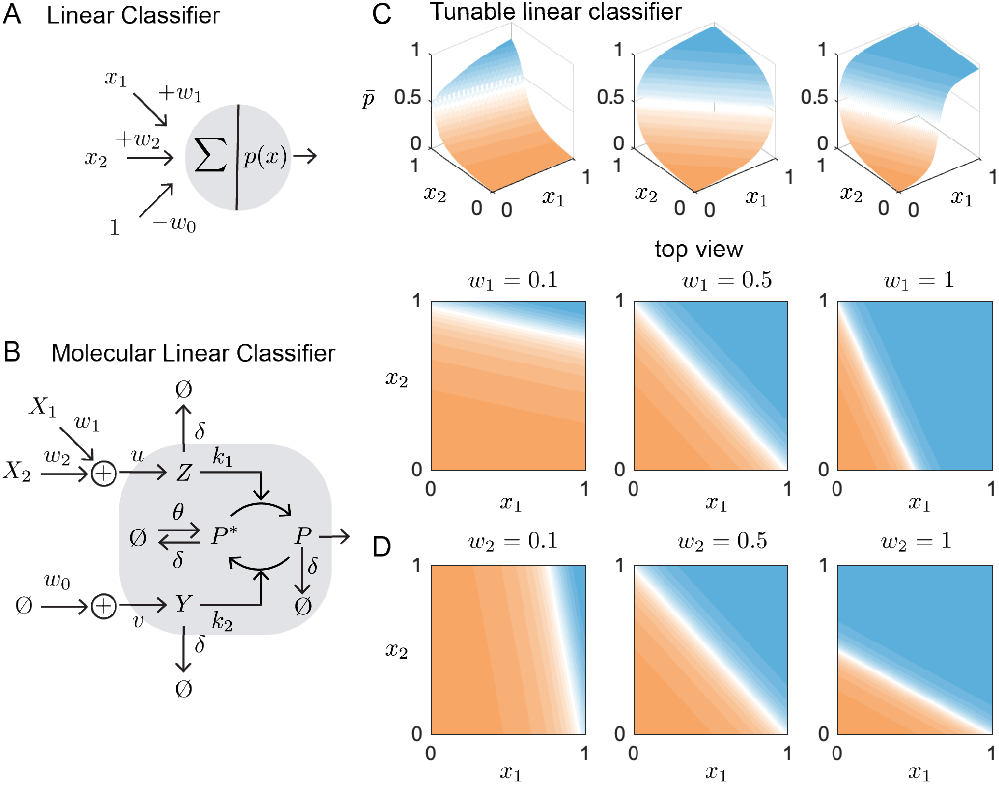
Three-input phosphorylation-based linear classifier with two positive weights and one negative weight. A) Design schematic of a single perceptron with three inputs presenting two positive and a single negative weight. B) Phosphorylation-based realization of the perceptron. C) Steady state solutions are shown as heat maps for a range of inputs *x*_1_ and *x*_2_. A top-view of the heat-maps for different values of weights *w*_1_, demonstrating the tunability of the decision boundary (white region). D) The effect of varying *w*_2_ changes the y-axis scale on the decision boundary (top-view heat-maps). Parameter for simulations *w*_0_ = *w*_1_ = *w*_2_ = 0.5, *θ* = *δ* = 1, *k*_1_ = *k*_2_ = 10, and *K*_1_ = *K*_2_ = 0.05.

Using analogous assumptions as in Section III to simplify the model leads to the ODE system:

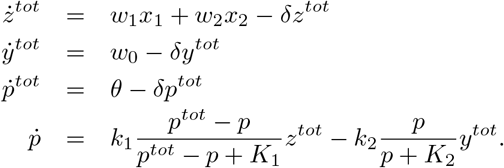

At steady state, 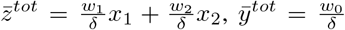, and 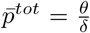. We can find 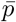 as a function of the system inputs by following the same procedure as in Section III, which eventually yields (25) where

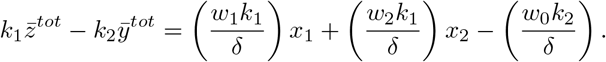

The terms inside the parenthesis are the effective weights of the system. Fig. 2C-D shows how the linear classification of the system changes when we change the production rates *w*_1_ and *w*_2_: Varying *w*_1_ rescales the x-axis, varying *w*_2_ rescales the y-axis, and varying *w*_0_ rigidly shifts the white region (not shown here).

### B. A network with one positive and two negative weights

As a second case study, we consider the design of a linear perceptron with one positive and two negative weights, as shown in Fig. 3A top. Hence, this reaction network features a single input *X*_2_, which produces *Z* with rate *w*_2_, and two inputs which produce *Y* – a constant production *w*_0_ and an input *X*_1_ associated with rate *w*_1_ – as shown in Fig. 3A (bottom). This leads to the ODE system:

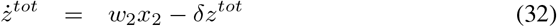

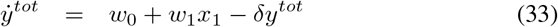

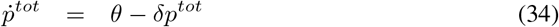

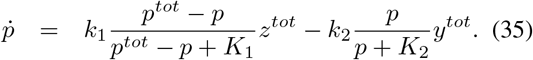

**Fig. 3.**
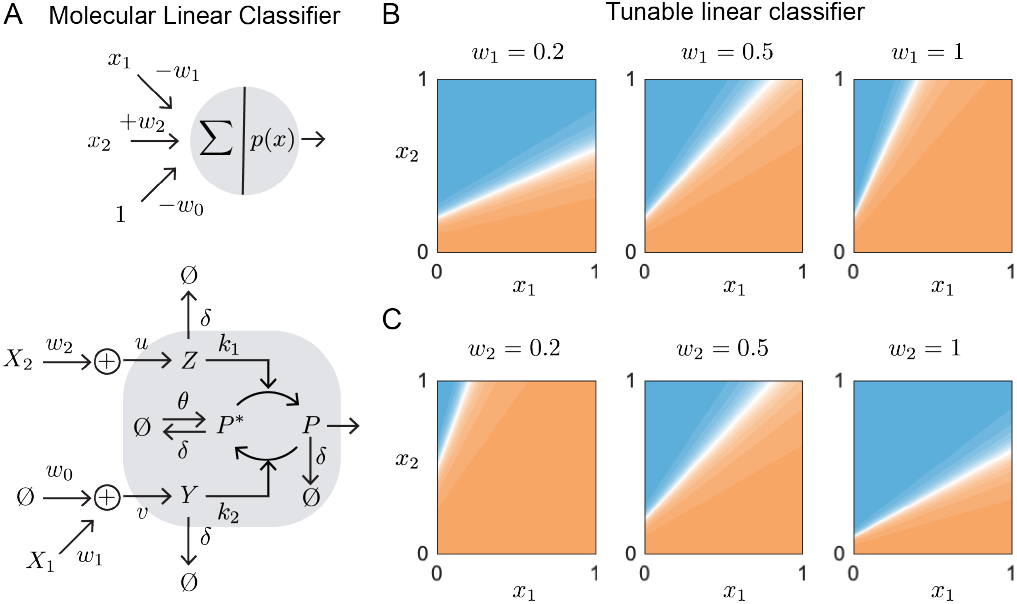
Three-input phosphorylation-based linear classifier with one positive and two negative weights. A) Design specification and molecular implementation of a single perceptron with three inputs, one positive and two negative weights. C) Steady state solutions for different values of weights *w*_1_, it results in rescaling the y-axis. D) Varying the production rate *w*_2_ rescale the x-axis. Parameter for simulations *w*_0_ = *w*_1_ = *w*_2_ = 0.5, *θ* = *δ* = 1, *k*_1_ = *k*_2_ = 10, and *K*_1_ = *K*_2_ = 0.05.

At the steady state, 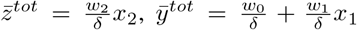, and 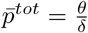, while 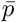 is expressed by (25) with

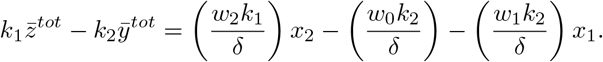

Again, varying the production rates of the system has an equivalent effect on the network’s linear classification: *w*_1_ and *w*_2_ rescale the x- and y-axes, respectively (Fig. 3B-C) and *w*_0_ rigidly shifts the white region (not shown here).

In this example, we can observe the importance of incorporating negatively-weighted input values: Changing the sign of *w*_1_ from positive to negative reorients the decision boundary and affords this perceptron greater representation power than the previous example. Specifically, a linear classifier with only positive weights (excluding the bias term), like the perceptron in Fig. 2, is incapable of producing the classification in Fig. 3 without negative weights. In the following sections, we will also demonstrate how tuning positive and negative weights in molecular networks can enable us to engineer more complex decision boundaries when multiple nodes are interconnected.

## V. Multi-Layer Biomolecular Neural Networks

Having shown how post-translational modification cycles can operate as a single biomolecular perceptron, we now study networks obtained by cascading many such perceptrons into a multi-layer system. In the following, we denote as *Biomolecular Neural Network* (BNN) a network including multiple perceptrons, that are organized into layers linked only by feedforward connections (cf. Fig. 4). Each perceptron in a layer processes input species *x*_*i,j*_ (*i* ∈ {1*, …, W*}, *j* ∈ 1*, …, D*) to produce output species *y*_*i,j*_, where *W* is the width and *D* is the depth of the network.

**Fig. 4.**
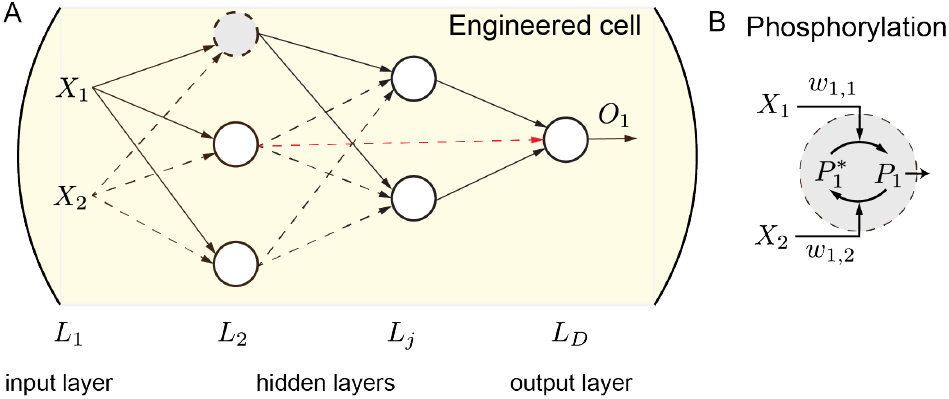
Biomolecular Neural Network. A) The schematic diagram for a BNN of depth *D* = 4 with two input species, *X*_1_ and *X*_2_, six biomolecular perceptrons in cicles, and one output species *O*_1_. Black arrows represent feedforward connections between neighboring layers, and red lines, feedforward connections spanning more than one layer. B) A simple biological implementation of each node based on phosphorylation reactions.

BNNs including layered perceptrons based on molecular sequestration have been studied in [4]. Here, we analyze the stability of the equilibria of a BNN composed of layered phosphorylation-dephosphorylation modules.

### Proposition 2

Consider a BNN with depth *D* and width *w*_*j*_ for each layer *j* (hence, *W* = Σ_*j*_ *w*_*j*_). Every equilibrium point is locally asymptotically stable.

*Proof:* The general Jacobian matrix for a feedforward

BNN network takes the lower block-triangular form:

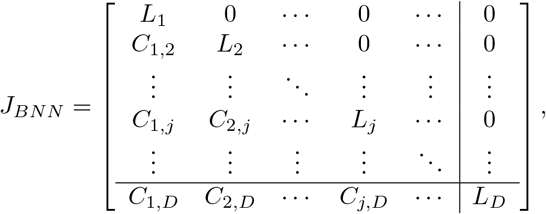

where

- matrix *C*_*j,k*_, with *j, k* ∈ {1*, …, D*} and *j < k*, describes how the variables in layer *j* affect the dynamics of the variables in layer *k*;
- matrix *L*_*j*_ represents the autonomous dynamics of layer *j* and has the block-diagonal form

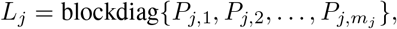

where *P*_*i,j*_ denotes the Jacobian matrix of the biomolecular perceptron in position *i, j*.

Since the Jacobian *J*_*BNN*_ is a block-triangular matrix, its characteristic polynomial *ψ*_*BNN*_(*s*) is the product of the characteristic polynomials of its diagonal blocks. Also, since each diagonal block *L*_*j*_ is block-diagonal, its characteristic polynomial is in turn the product of the characteristic polynomials *ψ*_*i,j*_(*s*) of the perceptrons *P*_*i,j*_ included in the corresponding layer. Therefore,

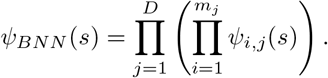

Since the generic phosphorylation-dephosphorylation perceptron is structurally asymptotically stable, as shown in Theorem 2, all the eigenvalues of matrix *J*_*BNN*_ have negative real part. Therefore, the BNN is asymptotically stable around any equilibrium.

## VI. Molecular Non-Linear Classifiers

In the previous section we demonstrated the stability of any feedforward BNN that includes a cascade of perceptrons based on post-translational modification cycles. In this section, we use three simulation examples to illustrate how feedforward BNNs may be used to build non-linear classifiers, highlighting their versatility and computational power.

### A. XNOR and XOR bio-molecular networks

Exclusive logical or (XOR) classification is a quintessential problem in the field of machine learning. It is the only logical operation which is not linearly separable, and has consequently motivated the use of multi-layer networks: Unlike single perceptrons, networks with two or more layers can perform non-linear classifications (e.g. XOR) provided appropriate weights.

For example, by adjusting its parameter values, the two-layer, three-node network shown in Fig. 5A can implement either XNOR or XOR logical functions. To conserve space, we do not show the detailed chemical reactions for this network, but describe the system with the following ODEs, following steps as given in previous sections:

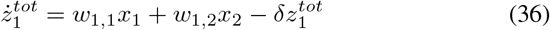

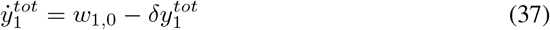

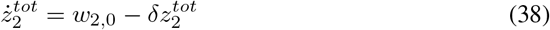

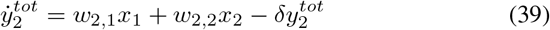

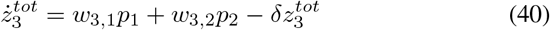

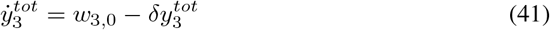

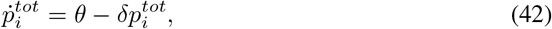

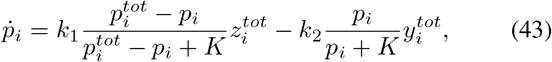

with *i* = 1, 2, 3.

**Fig. 5.**
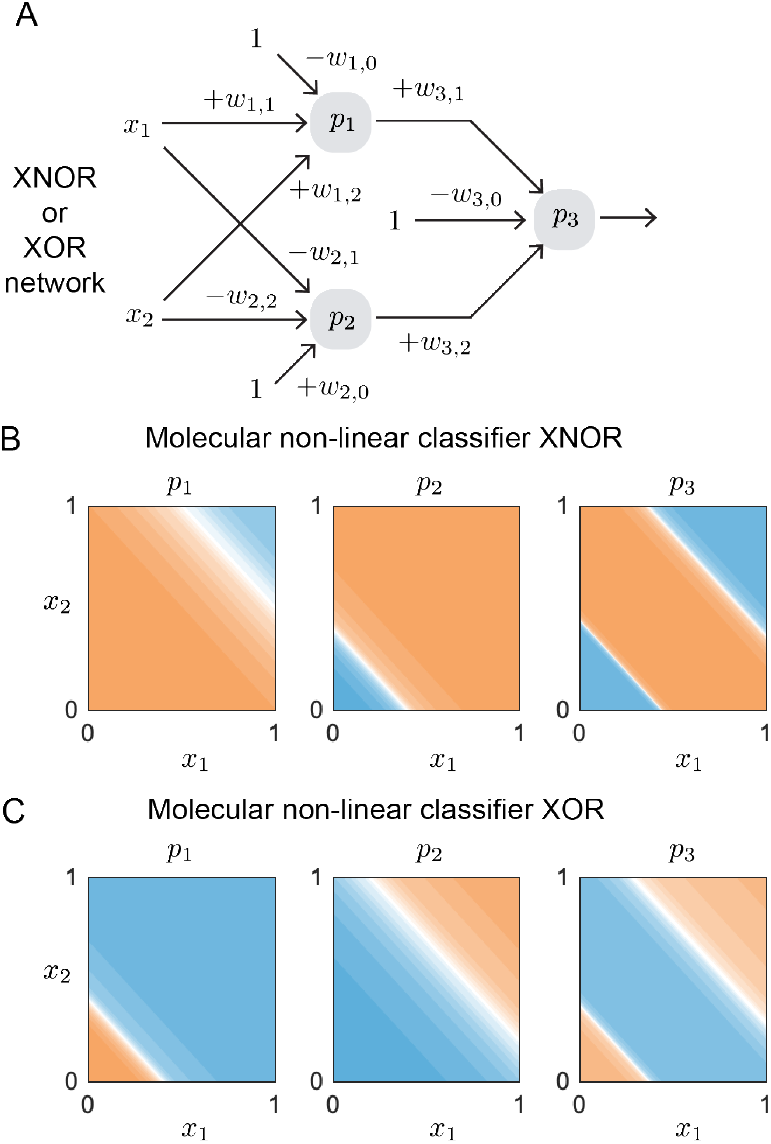
Phosphorylation-based XNOR and XOR classifiers. A) Network design. B) Steady state solution for an XNOR with optimized weights (*w*_1,0:2_ = [1.5, 1, 1], *w*_2,0:2_ = [0.4, 1, 1], *w*_3,0:2_ = [0.3, 1, 1]). C) Steady state solution for an XOR with optimized weights (*w*_1,0:2_ = [0.4, 1, 1], *w*_2,0:2_ = [1.2, 1, 1], *w*_3,0:2_ = [1.3, 1, 1]). The rest of the parameter for both simulations are *θ* = *δ* = 1, *k*_1_ = *k*_2_ = 10, and *K* = 0.05.

Notably, the network topology above may carry out either classification by only modifying the bias weights. In Fig. 5, we display the intermediate transfer functions given by perceptrons *p*_1_ and *p*_2_ (Fig. 5B and C, Left and Middle) for each non-linear classifier. To obtain an XNOR classification, we require *p*_1_ and *p*_2_ to have oppositely signed weights applied to *x*_1_ and *x*_2_ – positive weights for *p*_1_ and negative for *p*_2_ – and a larger magnitude of bias for *w*_1,0_ than *w*_2,0_ (see Fig. 5B). Under these conditions, the entire network represents an XNOR function; the output of *p*_3_ is maximal when *x*_1_ and *x*_2_ both take on low or high values. In contrast, if we decrease the magnitude of bias *w*_1,0_ and increase biases *w*_2,0_ and *w*_3,0_, the same network topology can perform an XOR operation, which has the opposite output (see Fig. 5C).

This case study illustrates an important general property of BNNs: By changing the weights of each node, the classification boundary, represented by the white area of each plot, can be adjusted in a predictable way and to represent highly dissimilar decision-making functions. Biologically, such changes are tantamount to adjustments of the rate parameters of the network’s constituent reactions; no modifications to its underlying topology are required.

### B. More complex decision-making

Although the previous classifier is able to deliver a non-linear function of its inputs, its network topology is incapable of creating a closed decision boundary. To achieve this form of classification, we add a perceptron to the first layer of the network, creating a four-node network (Fig. 6). We then optimize the weights of 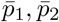 and 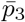 to enclose a central region of the classification space, as shown in Fig. 6B. Only where their maximal values (i.e. blue regions) overlap is the threshold value of the fourth node exceeded. This "activated" region forms a closed boundary where the output of the entire network is high. Moreover, by modifying the weights of each node it is possible to shift, scale, and rotate this closed boundary and enclose alternative regions (not shown here). The ODEs describing this network are given below:

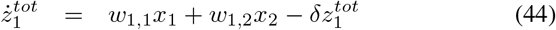

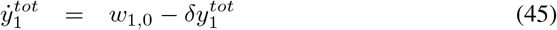

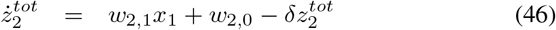

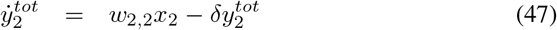

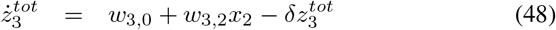

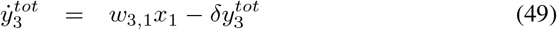

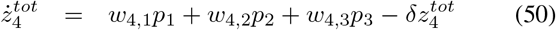

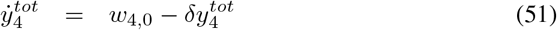

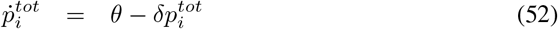

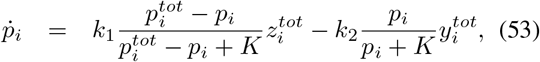

for *i* = 1, 2, 3, 4.

**Fig. 6.**
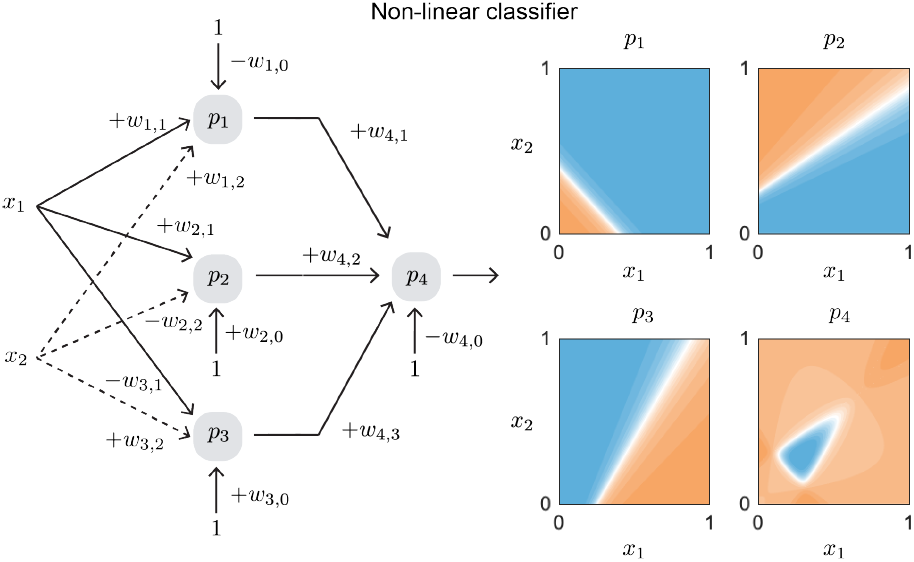
Phosphorylation-based non-linear classifiers. Network design for a closed-type decision making with four nodes. Parameter for simulations *w*_1,0:2_ = [0.2, 0.5, 0.5], *w*_2,0:2_ = [0.2, 0.5, 0.8], *w*_3,0:2_ = [0.2, 0.8, 0.5], *w*_4,0:3_ = [1.3, 0.5, 0.5, 0.5] *θ* = *δ* = 1, *k*_1_ = *k*_2_ = 10, and *K* = 0.05.

## VII. Discussion

We have discussed how, under some assumptions, a broad class of biomolecular signaling networks have the capacity to operate as molecular perceptrons, due to their sigmoidal input-output behavior with a tunable threshold, that can integrate additively the contribution of multiple inputs. In particular we have shown that the perceptron model arising from this type of network includes inputs that are weighted both positively and negatively. A great advantage of realizing and tuning negative weights is that they make it possible to optimize networks for classification. We illustrated this concept by simulating linear and non-linear classifiers based on phosphorylation/dephosphorylation cycles. This manuscript extends to signaling networks our earlier work that demonstrated how molecular sequestration motifs have the capacity to implement biomolecular neural networks [4]. While here we focus exclusively on demonstrating the computational and classification power of signaling networks, our goal for future work is to build on this approach to examine the contribution of cross-talk to the decision boundary of post-translational modification cycles.

Our analysis complements theoretical and experimental efforts toward building complex molecular networks such as chemical Boltzmann machines [7] and DNA-based classifiers [13], [11]. As molecular neural networks are becoming relevant for applications that include *in vitro* diagnostics [12] and *in vivo* cellular classifiers[13], we expect that methods to harness the computational power and implementation options of BNNs will make it possible to expand the ways we can engineer living cells for advanced decision making.

## VIII. ACKNOWLEDGMENTS

EF and CCS are sponsored by NSF/BBSRC award 2020039. The content of the information does not necessarily reflect the position or the policy of the Government, and no official endorsement should be inferred.

## Notes

### Competing Interest Statement

The authors have declared no competing interest.

